# Prolonged exposure to hypergravity increases number and size of cells and enhances lignin deposition in the stem of *Arabidopsis thaliana* (L.) Heynh

**DOI:** 10.1101/2023.06.15.545094

**Authors:** Hironori Shinohara, Masaki Muramoto, Daisuke Tamaoki, Hiroyuki Kamachi, Hiroshi Inoue, Atsushi Kume, Ichirou Karahara

## Abstract

We have performed a lab-based hypergravity cultivation experiment using a centrifuge equipped with a lighting system and examined long-term effects of hypergravity on the development of the main axis (stem) of the Arabidopsis (*Arabidopsis thaliana* (L.) Heynh.) primary inflorescence. Plants grown under 1 × *g* (gravitational acceleration on Earth) conditions for 20-23 days and having the first visible flower bud were exposed to hypergravity at 8 × *g* for 10 days. We analyzed the effect of prolonged hypergravity conditions on growth, lignin deposition, and tissue anatomy of the main axis. As a result, the length of the main axis decreased and cross-sectional area, dry mass per unit length, cell number, lignin content of the main axis significantly increased under hypergravity. Lignin content in the rosette leaves also increased when they were exposed to hypergravity during their development. Except for interfascicular fibers, cross-sectional areas of the tissues composing the internode significantly increased under hypergravity in most type of the tissues in the basal part than the apical part of the main axis, indicating that the effect of hypergravity is more pronounced in the basal part than the apical part. The number of cells in fascicular cambium and xylem significantly increased under hypergravity both in the apical and basal internodes of the main axis, indicating a possibility that hypergravity stimulates procambium activity to produce xylem element more than phloem element. The main axis was suggested to be strengthened through changes in its morphological characteristics as well as lignin deposition under prolonged hypergravity conditions.

## Introduction

Development of vascular tissues, having intensively lignified secondary cell walls, has been postulated to be essential for the evolution of land plants for both physical and physiological reasons, i.e., for resistance to 1 × *g* gravity and for adaptation to the dry environment on land. Lignin in the secondary cell wall imparts rigidity to the plant bodies. According to previous space experiments, lignin formation is suppressed under microgravity conditions in space. In addition, hypergravity experiments performed on the ground also revealed that exposure of Arabidopsis peduncles to 300 × *g* for 24 h enhanced primary xylem development (Nakabayashi et al. 2006), promoted formation of secondary cell walls and deposition of acetyl bromide-soluble lignin (ABS-lignin) (Tamaoki et al. 2006), and upregulated expressions of genes related to lignin synthesis (Tamaoki et al. 2009). The content of ABS-lignin increased gradually from the apical to the basal regions of azuki bean (*Vigna angularis* Ohwi et Ohashi) epicotyls when they were treated with 300 × *g* for 6 h (Wakabayashi et al. 2009). Therefore, lignin formation is suggested to play a role in gravity resistance mechanism (Soga 2013). On the other hand, no significant promotive effect was observed in ABS-lignin content in the leaf of *Triticum aestivum* L. grown under microgravity for 21 d (Stutte et al. 2006) as well as in the stem of *Brassica rapa* L. ‘Wisconsin Fast Plant’ grown under 4 × *g* for 16 d (Allen et al. 2009). The issue whether lignin formation is regulated in response to gravity in plants is still controversial and, therefore, effects of prolonged hypergravity exposure on lignin deposition needs to be further examined in Arabidopsis.

Different from the effect of hypergravity on lignin content, however, regarding tissue morphology, hypergravity treatment increased the size of cells significantly in the pith, cortex, and vascular tissue in the stem of *B. rapa* L. even though it did not affect lignin content (Allen et al. 2009). Although most of previous studies using hypergravity have low exposure durations (< 5 h) (Hosamani et al. 2022), Allen et al. (2009) employed a large-diameter centrifuge available at a space center. Such a facility provides an ideal environment for a long-term hypergravity experiment but its availability is limited. Instead, a custom-built centrifuge equipped with a lighting system is developed and being employed for long-term hypergravity experiment, and gravity resistance responses of *Physcomitrium* (*Physcomitrella*) *patens* (Hedw.) Bruch et Schimp., such as, decrease in shoot axial growth and increase in shoot diameter, are demonstrated (Takemura et al. 2017a; Takemura et al. 2017b). In the present study, we also successfully designed a lab-based prolonged hypergravity experiment using a centrifuge equipped with a lighting system, analyzed the effect of exposure to 8 × *g* for 10 days on lignin deposition in the peduncle, and on tissue anatomy comparing it between the positions near the apex and the base of the primary inflorescence to discuss the effects of hypergravity at the cellular level, and discussed comparing the results with those of our previous study obtained using hypergravity conditions of 300 × *g* for 1 d.

## Materials and Methods

### Plant materials and hypergravity treatment

Plants of Arabidopsis (*Arabidopsis thaliana* (L.) Heynh.) of Columbia-0 line were used for the experiments. After surface sterilization with 99 % (v/v) ethanol for 10 s, 4 seeds were planted on 30 mL of 1.0 % (w/v) agar containing Murashige and Skoog medium (Wako Pure Chemical, Tokyo, Japan) in a polycarbonate container (Plant Box, Biomedical Science, Tokyo, Japan; 80 mm in diameter, 11.5 mm in height), and kept at 4 °C for 3 d, and then allowed to grow at 23 °C for 10 d for analysis of rosette leaves or for 20-23 d for analysis of the primary inflorescences under 1 × *g* under continuous white light provided with a bank of fluorescent tubes (Mellow white 20 W daylight type; Toshiba Lighting & Technology Co., Yokosuka, Japan) with the intensity being 130 μmol m^2^ s^-1^ at plant level. For analysis of growth of the primary inflorescence, 20-23-d-old plants having the primary inflorescence of 10 mm in length (Arabidopsis growth stage number 5.10, the first flower bud visible; Boyes et al. 2001) were selected. For analysis of rosette leaves, 10-d-old plants having 2 visible rosette leaves which are larger than 1 mm in length (Arabidopsis growth stage number 1.02) (Boyes et al. 2001) were selected. These plants were exposed to hypergravity at 8 × *g* for 10 d at 25 °C using a centrifuge (Model 8420, Kubota Corporation Co., Tokyo, Japan; Effective radius, 185 mm) equipped with fluorescent bulbs (Neoball Z EFD21ED 21 W, Toshiba Lighting & Technology Co.), where its intensity being 80 μmol m^2^ s^-1^. For 1 × *g* control, the containers were placed at 25 °C under the fluorescent bulbs without centrifugation.

### Measurements of morphology and dry mass of peduncles and rosette leaves

After measurement of the length of the main axis of the primary inflorescence, which is composed of rachis and peduncle, the main axis and rosette leaves were cut and dried at 60 °C for 12 h. Their dry mass was measured using an ultramicrobalance (SE2, Sartorius Japan, Tokyo, Japan).

### Fluorescence microscopy

Fresh transverse sections of 100-μm thickness were cut from the lowermost internode just above the last rosette leaf, i.e., peduncle (referred to as the basal position), internode at middle position, and that near the apical position of the main axis of the primary inflorescence exposed to 1 × *g* or 8 × *g* using a vibrating microtome (Linear slicer Pro 7, Dosaka EM, Kyoto, Japan) and observed under a fluorescence microscope (BX-50 FLA, Olympus, Tokyo, Japan) equipped with a filter assembly for excitation by ultraviolet light (U-MWU: excitation filter, BP330-385; absorption filter, BA420; dichroic mirror, DM-400; Olympus). Fluorescence digital micrographs were taken using a digital camera (Coolsnap cf, Nippon Roper, Tokyo, Japan) fitted to the microscope.

### Lignin quantification

For lignin quantification in the main axis of the primary inflorescence, 20-23-d-old plants having the primary inflorescence of 10 mm in length were treated with 8 × *g* for 10 d. For lignin quantification in rosette leaves, 20-23-d-old plants having the primary inflorescence of 10 mm in length or 10-d-old plants were treated with 8 × *g* for 10 d. Each main axis or rosette leaf was dried overnight at 60 °C. After measuring its dry mass, each organ was cut into pieces with a razor blade. The quantification of ABS-lignin was carried out according to the methods described previously (Morrison 1972, De Jaegher et al. 1985).

### Embedding specimens and quantitative anatomical analyses

Five-mm segments cut from an apical and a middle part, which are in the internodes of the rachis, and a basal part, which is in the internode of the peduncle, of the main axis of the primary inflorescence of plants, which were treated with 1 × *g* or 8 × *g* for 10 d after grown under 1 × *g* for 20-23 d, were fixed in FAA solution containing 5 % (v/v) formalin, 5 % (v/v) acetic acid, and 50 % (v/v) ethanol for 12 h. The fixed segments were washed, dehydrated by passage through a graded ethanol and propylene oxide series, and embedded in Quetol resin (Nisshin EM, Tokyo, Japan). Transverse semi-thin sections of 1-μm thickness were cut using ultramicrotome, observed under a bright field or a fluorescence microscope as described above. Digital micrographs were taken as described above.

### Statistical analysis

For testing statistical difference between two samples, normality and variance equality of population distribution was examined. When both of the assumptions were hold, Student’s *t*-test was performed. When neither of the assumptions were hold, Wilcoxon rank sum test was performed. When normality was hold but variance equality was not, Welch’s *t*-test was performed instead.

## Results

### Effects of prolonged hypergravity on the growth of primary inflorescence

Effects of long-term exposure to hypergravity on the morphology of Arabidopsis plants were examined. The primary inflorescence could continue to grow even under 8 × *g* (Fig. 1a). However, the primary inflorescence could not keep standing in an upright position under 8 × *g* while it could under 1 × *g* (Fig. 1b, c). While the lower part of the primary inflorescence was laid down or slanted, its upper part could grow upwards with the support of the inner wall of the cylindrical container. Time-course examination of the length of the primary inflorescence revealed that the growth of the primary inflorescence was significantly suppressed under 8 × *g* from 8 days after the start of the treatment (Fig. 1a).

**Fig. 1.**
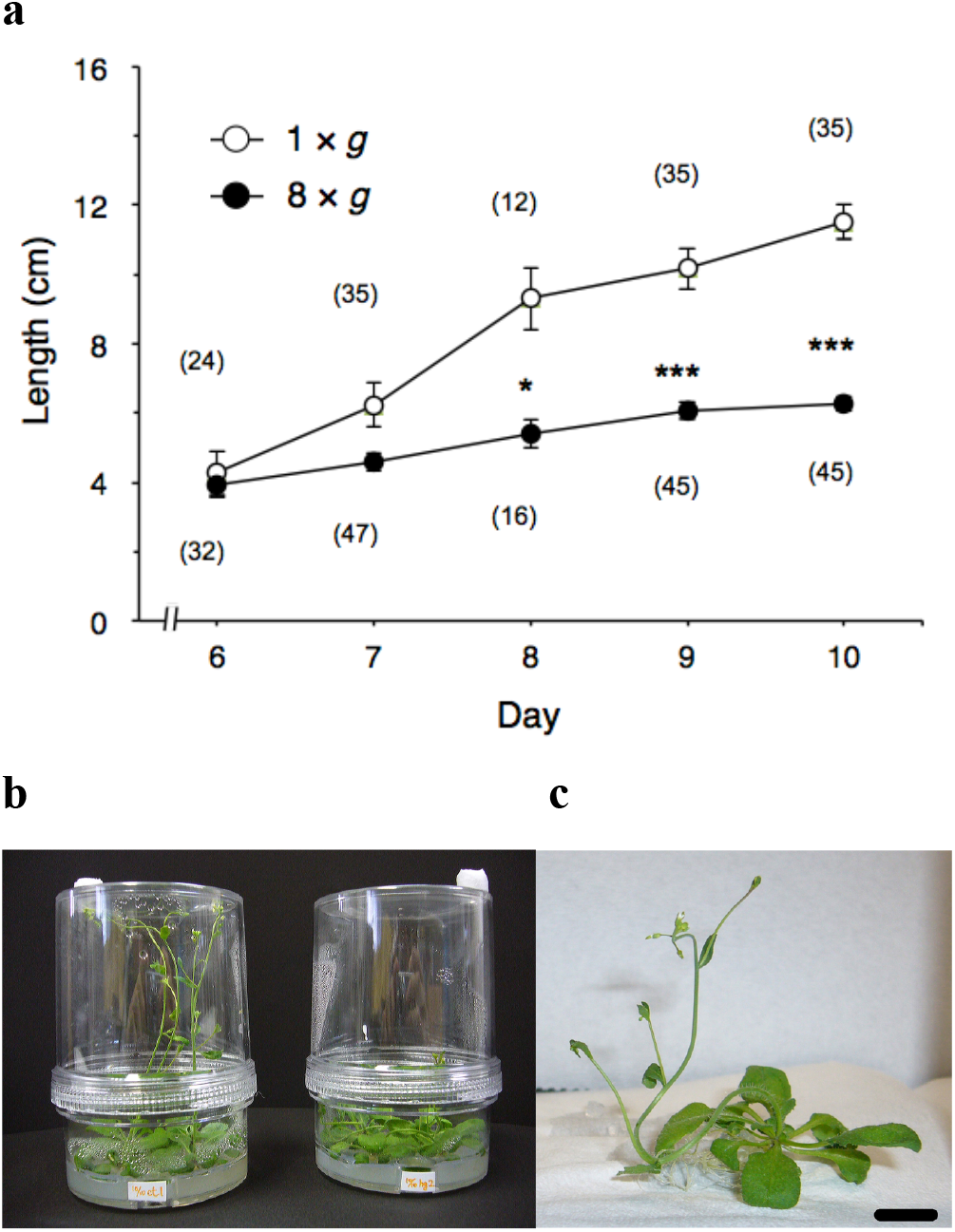
Effect of 8 × *g* hypergravity on the growth of the main axis of the primary inflorescence (**a**) and the morphology of the plants (**b, c**) of Arabidopsis. **a** Time-course change in the length of the primary inflorescence plotted against days after the start of 8 × *g* hypergravity exposure. Values are mean ± SE. ***, *P* < 0.001; *, *P* < 0.05 (Wilcoxon rank sum test, two-tailed). The numbers of samples are shown in parenthesis. **b** Morphology of the plants exposed to 8 × *g* hypergravity (right) or with 1 × *g* (the control, left) for 10 days. **c** A close-up view of a typical plant exposed to 8 × *g*. Scale bar 1 cm

Length of the primary inflorescence significantly decreased and dry mass per unit length of the main axis of the primary inflorescence significantly increased after the 10-d exposure to 8 × *g* (Fig. 2a, b). Figure 3 shows that the first to fourth internode lengths of primary inflorescences were significantly shorter in the 8 × g treatment compared to the 1 × g treatment.

**Fig. 2.**
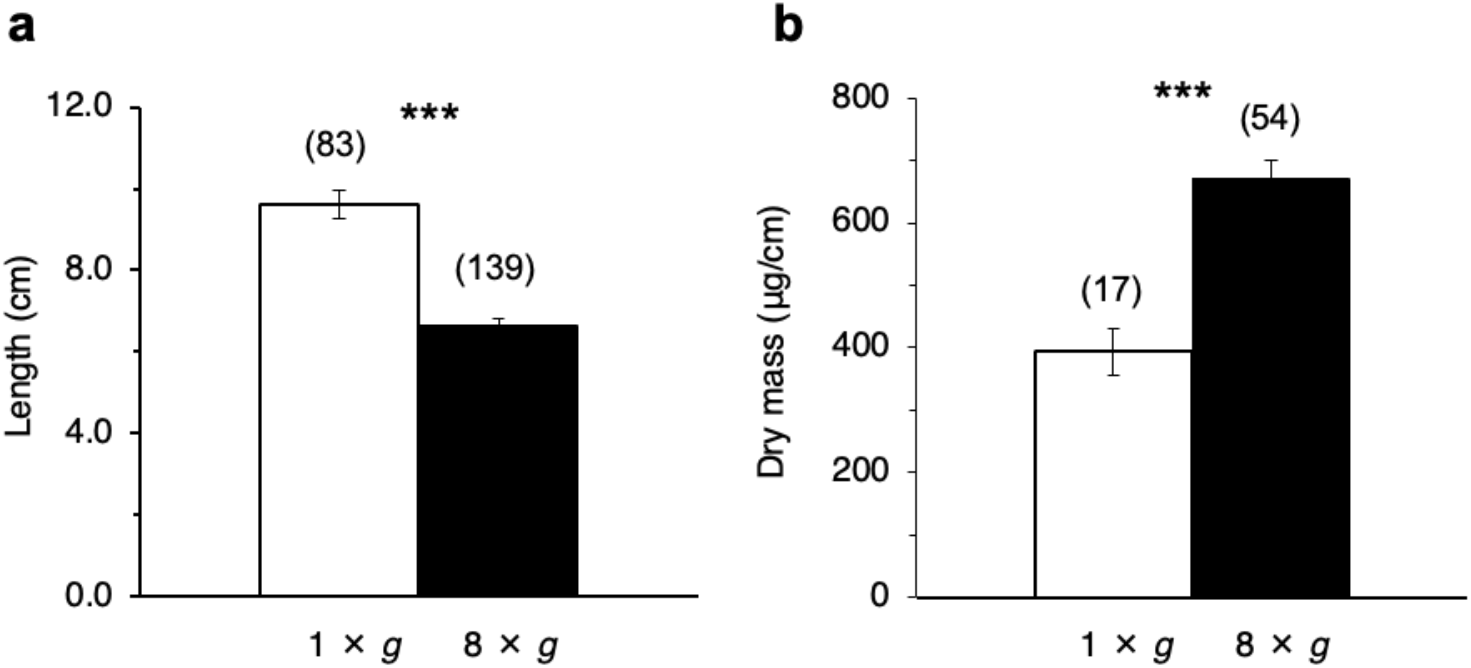
Effect of 10-d exposure to 8 × *g* hypergravity on the length (**a**) and dry mass per unit length (**b**) of the main axis of the Arabidopsis primary inflorescence. Values are presented as mean ± SE. Sample numbers are shown in parentheses. Statistical differences were tested using Wilcoxon rank sum test (two-tailed, ***, *P* < 0.001).

**Fig. 3.**
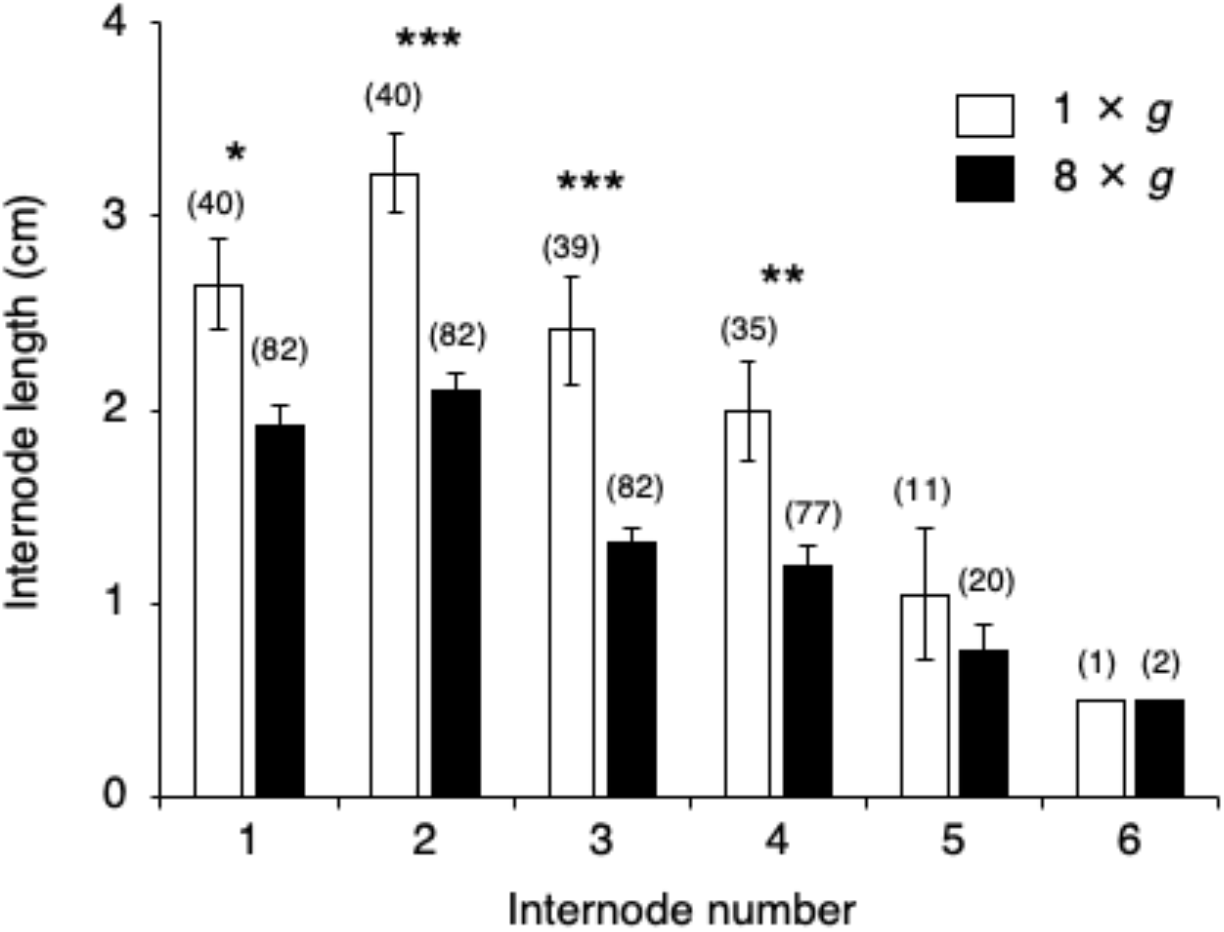
Effect of 10-d exposure to 8 × *g* hypergravity on internode lengths of the main axis of the Arabidopsis primary inflorescence. The first internode is defined as that located just above the uppermost rosette leaf. Values are presented as mean ± SE. Sample numbers are shown in parentheses. Statistical differences were tested within the same internode number using Wilcoxon rank sum test (two-tailed, ***, *P* < 0.001; **, *P* < 0.01; *, *P* < 0.05).

### Effects of prolonged hypergravity on the morphology of xylem and lignin content

We performed anatomical observation of the main axis to examine whether lignin deposition is enhanced by hypergravity exposure. Figure 4 shows that primary xylem recognized by blue autofluorescence of lignin developed from the apical to the basal position of the main axis either under 1 × *g* or 8 × *g* condition. While only xylem was observed either at the apical and middle position of the internode of the primary inflorescence of approximately one-month old plants (Fig. 4a, b, c, d; Fig. 5a), interfascicular fibers were observed besides xylem at the basal position (Fig. 4e, f, Fig. 5c, d). Nevertheless, secondary xylem development due to vascular cambium activity has not yet been observed. Interestingly, autofluorescence in xylem and interfascicular fibers due to lignin deposition appears stronger at each position in the main axis of the primary inflorescence exposed to hypergravity (Fig. 4b, d, f).

**Fig. 4.**
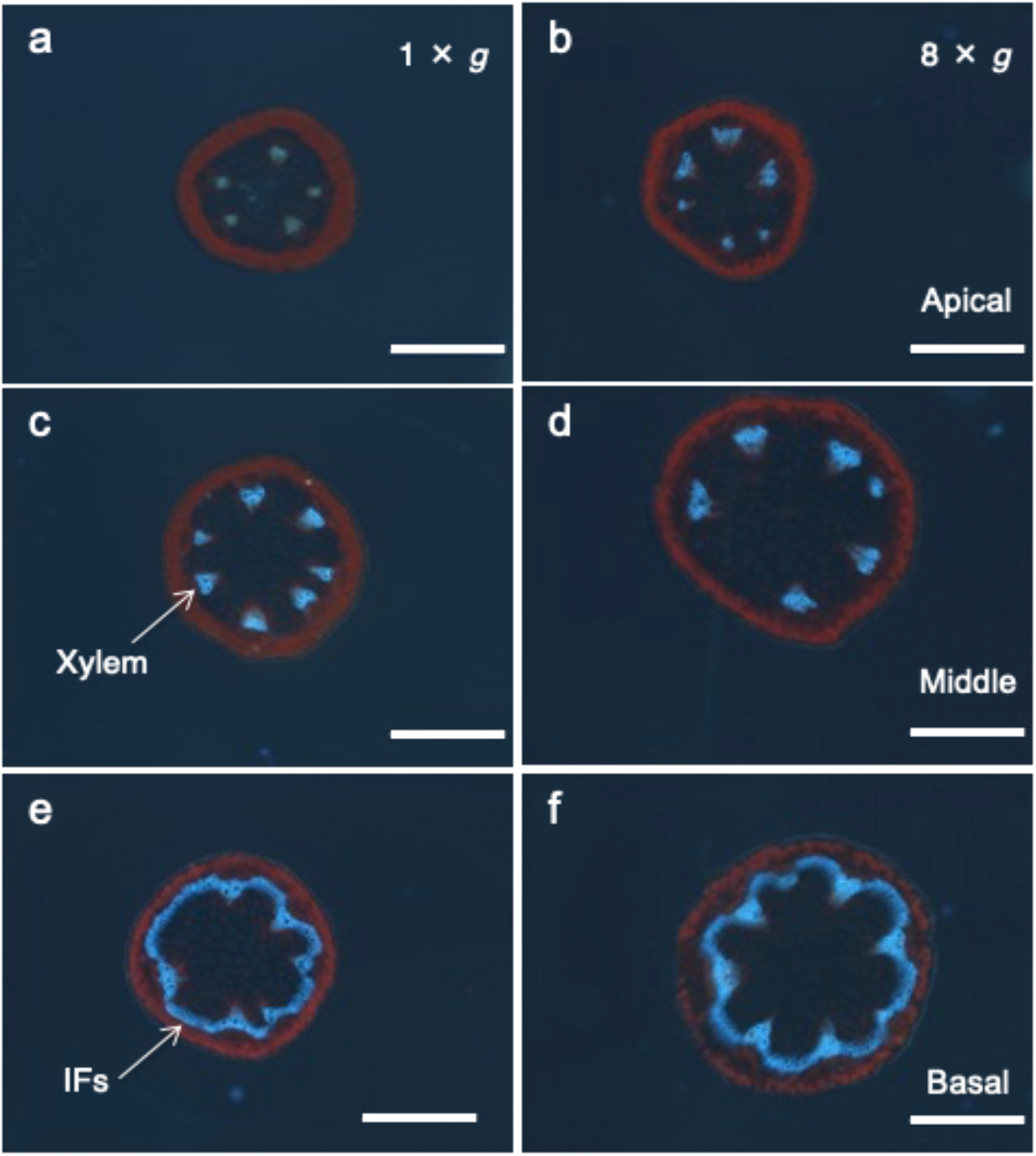
Effect of 10-d exposure to 8 × *g* hypergravity on autofluorescence of xylem vessels in internodes of the main axis of the Arabidopsis primary inflorescence. Fluorescence micrographs of transverse sections were obtained from the apical (**a, b**), middle (**c, d**), or basal positions (**e, f**) in the main axis of the primary inflorescence of a typical plant exposed to 1 × *g* (**a, c, e**) or 8 × *g* hypergravity (**b, d, f**). Red fluorescence shows autofluorescence of chloroplasts and blue fluorescence, which is due to lignin deposition, shows autofluorescence of xylem and interfascicular fibers. IFs: interfacicular fibers. Scale bars 0.5 mm

**Fig. 5.**
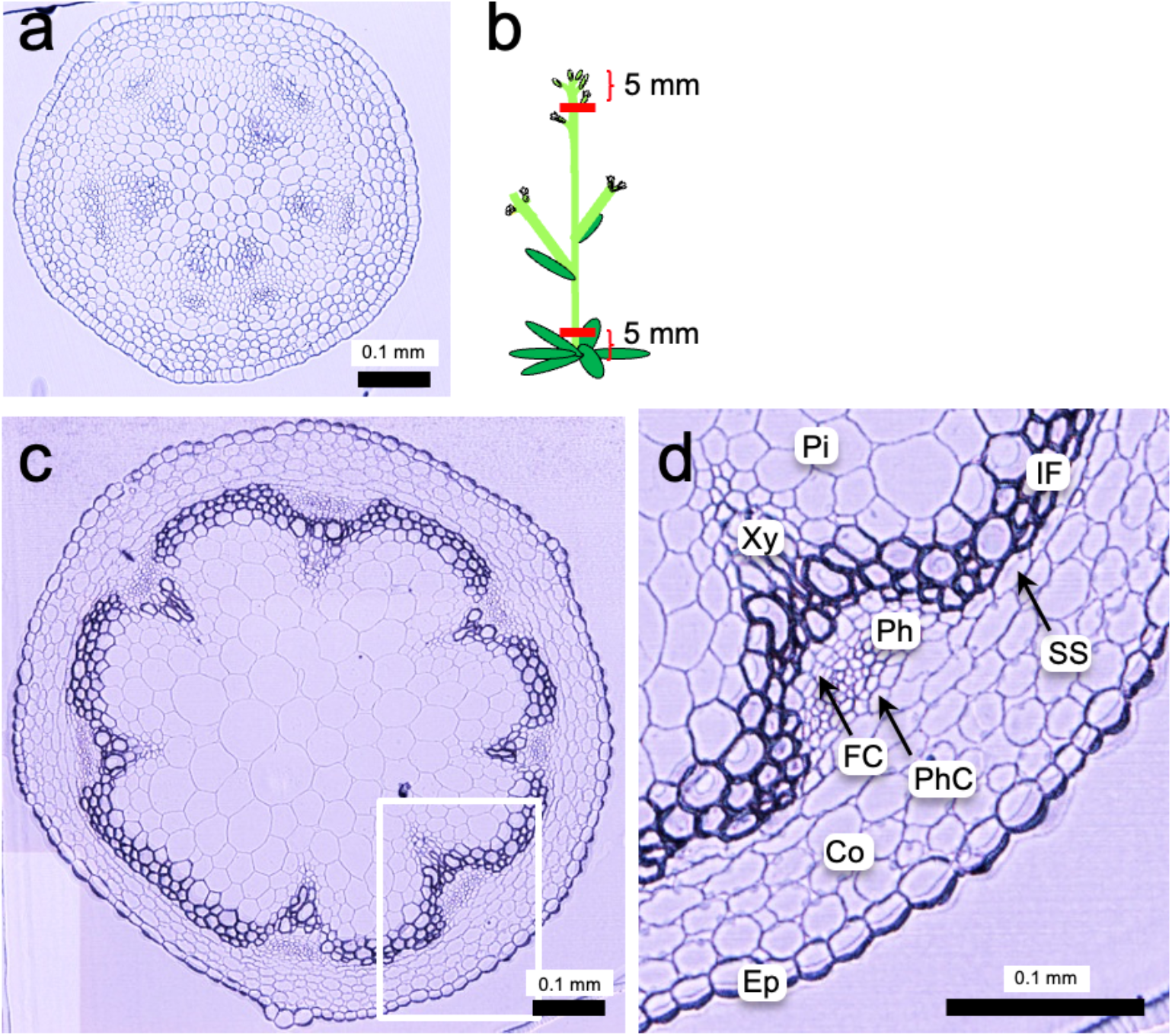
Anatomical observation of tissues composing the internodes of the main axis of the Arabidopsis primary inflorescence. Light micrographs of cross sections cut at 5 mm below the shoot apex (**a**) or at 5 mm above the base (**c**) of the main axis grown under 1 × *g* for a month (the control). (**b**) Schematic illustration showing the morphology of an Arabidopsis plant and the positions where cross sections were cut. (**d**) Higher magnification view of the area shown in a white rectangular in (**c**). Ep, epidermis; Co, cortex; SS, starch sheath; PhC, phloem cap; Ph, phloem; FC, fascicular cambium; IF, interfascicular fiber; Xy, xylem; Pi, pith. Scale bars 0.1 mm

To test whether lignification is also promoted under prolonged hypergravity in Arabidopsis, ABS-lignin was quantified for the main axis of the primary inflorescence as well as the rosette leaves of plants exposed to 8 × *g* for 10 d. As a result, lignin content significantly increased due to prolonged hypergravity exposure in the main axis of the primary inflorescence (Fig. 6a) while not significantly in the rosette leaves when they were exposed after they had been grown under 1 × *g* for 20-23 d and were at the Stage 5.10 (Fig. 6b). Interestingly, however, lignin content significantly increased under 8 × *g* in the rosette leaves when they were exposed after they had been grown under 1 × *g* for 10 d and were at the Stage 1.02 (Fig. 6c).

**Fig. 6.**
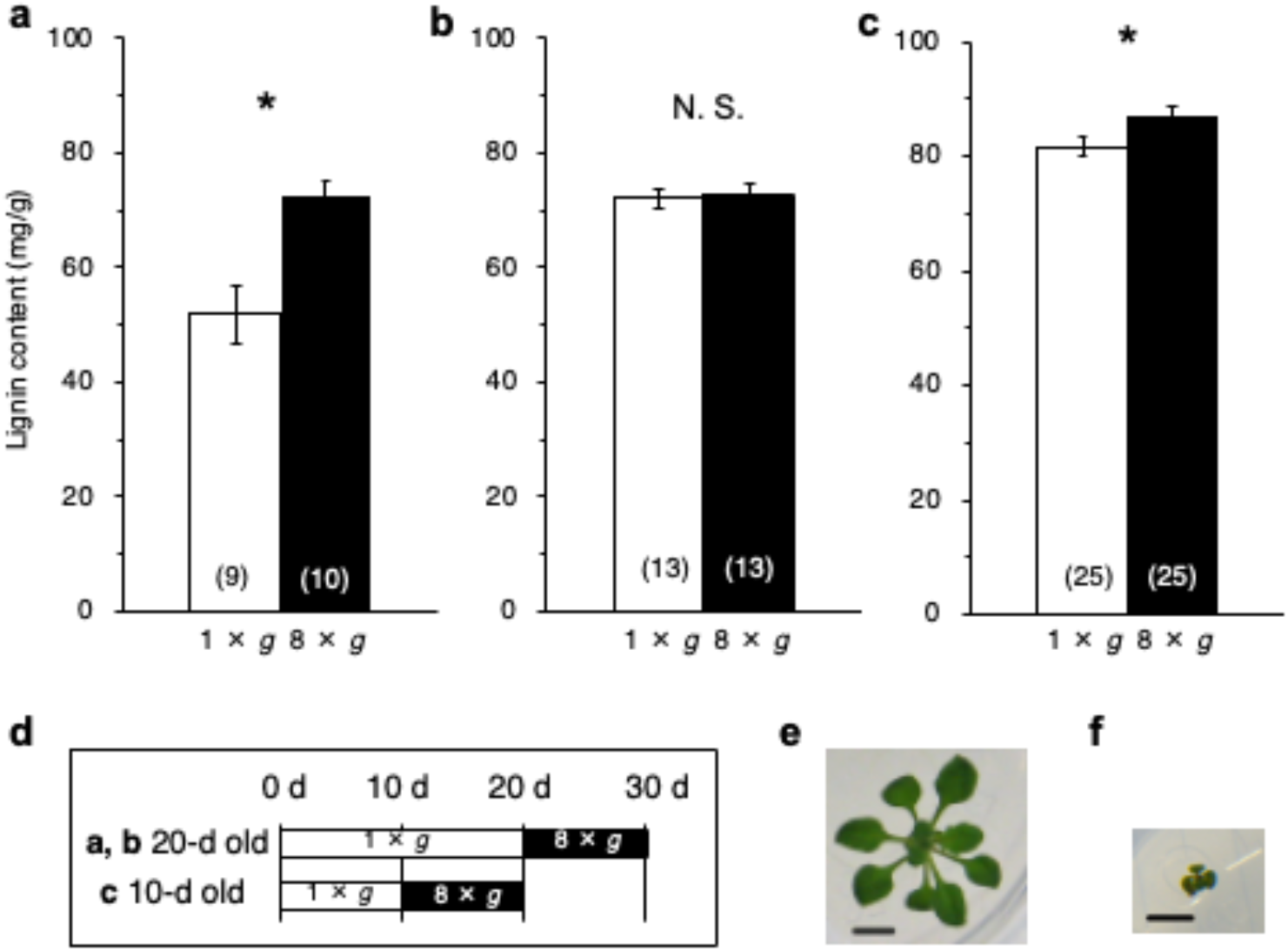
Effects of 10-d exposure to 8 × *g* hypergravity on the content of *ABS-lignin* (acetyl bromide-soluble lignin) in the main axis of the primary inflorescence (**a**) or the rosette leaves (**b, c**) per an Arabidopsis plant. Lignin quantification of the main axis of the primary inflorescence (**a**) and the rosette leaves (**b**) was first performed using plants treated with 8 × *g* hypergravity for 10 days after grown under 1 × *g* for 20-23 days (Stage 5.10, first flower buds visible, (Boyes et al. 2001)). Lignin quantification of the rosette leaves was also performed using plants treated with 8 × *g* hypergravity for 10 days after grown under 1 × *g* for only 10 days (Stage 1.02, 2 rosette leaves with their size > 1 mm, (Boyes et al. 2001)) (**c**). Statistical differences were tested using Wilcoxon rank sum test (two-tailed, *, *P* < 0.05). (**d**) A schematic diagram showing a difference in the timing of gravity treatments. A top view photograph of a typical plant grown under 1 × *g* for 20-23 days (Stage 5.10) **(e)** or for 10 days (Stage 1.02) (**f**), taken before the hypergravity exposure. Scale bars 5 mm

### Effects of prolonged hypergravity on the morphology of tissues composing the main axis

Quantitative anatomical analysis was performed to examine effects of prolonged hypergravity on the morphology of tissues composing the internode of the primary inflorescence at cellular level (Fig. 5). Figure 7 shows effects of 10-d exposure to 8 × g on the cross-sectional area, cell number, and cell size, which were calculated by dividing the cross-sectional area of a tissue by the cell number, of tissues composing the internode of the Arabidopsis primary inflorescence at the position 5 mm below the shoot apex or from the position 5 mm above the base of the main axis.

**Fig. 7.**
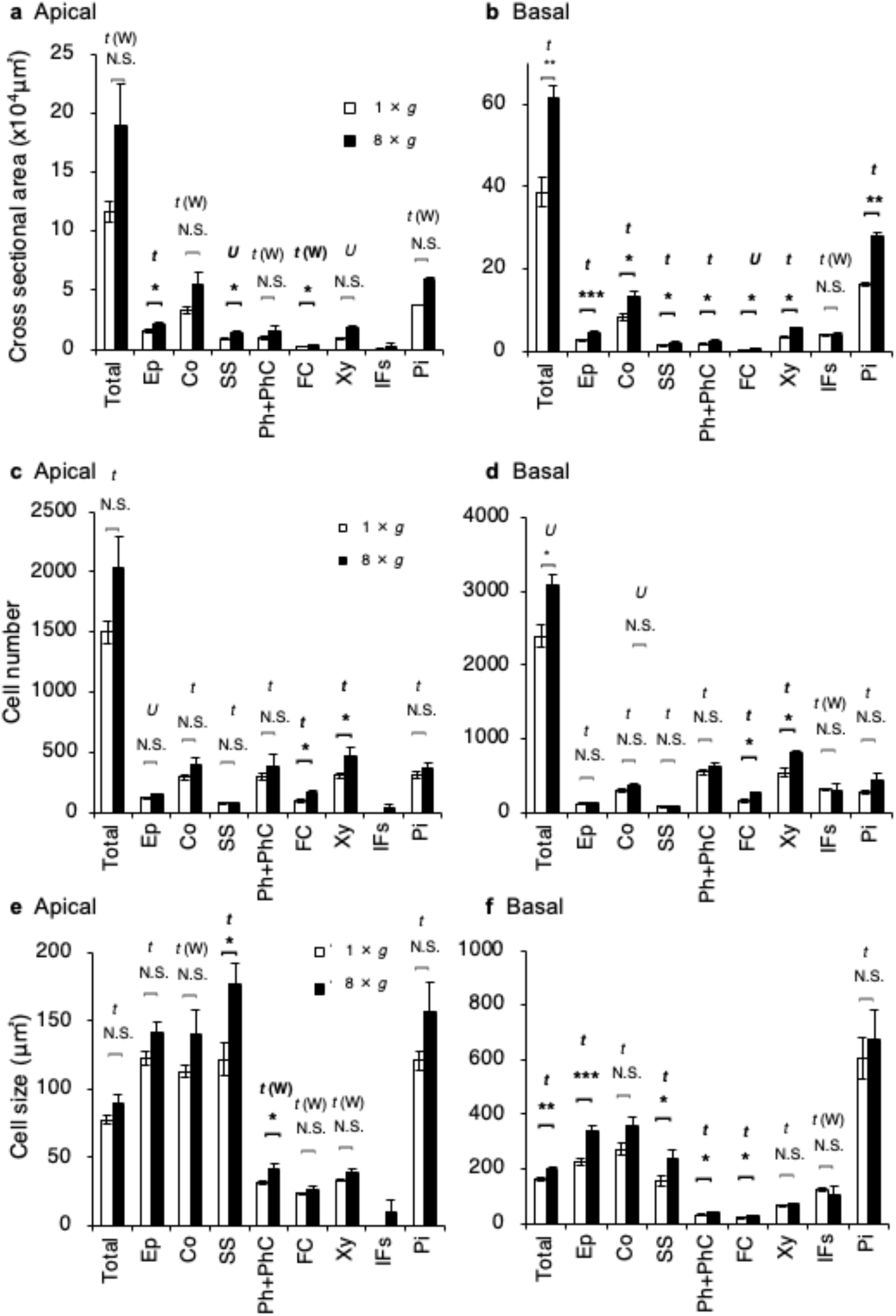
Effects of 10-d exposure to 8 × *g* hypergravity on the cross-sectional area (**a, b**), cell number (**c, d**), or cell size (**e, f**) of tissues composing the main axis of the Arabidopsis primary inflorescence at the position 5 mm below the shoot apex (**a, c, e**) or from the position 5 mm above the base of the peduncle (**b, d, f**). Ep, epidermis; Co, cortex; SS, starch sheath; Ph, phloem; PhC, phloem cap; FC, fascicular cambium; Xy, xylem; IFs, interfascicular fibers; Pi, pith. Statistical differences were tested within the same tissue using Student’s *t*-test (*t*), Welch’s *t*-test (*t*(W)), or Wilcoxon rank sum test (*U*) (two-tailed, n = 5). N.S., Not Significant; ***, *P* < 0.001; **, *P* < 0.01; *, *P* < 0.05.

As a result, cross-sectional area significantly increased due to prolonged exposure to hypergravity in most type of the tissues in the basal part except for interfascicular fibers (IFs) (Fig. 7b) while that significantly increased only in the epidermis (Ep), starch sheath (SS), and fascicular cambium (FC) in the apical part (Fig. 7a). Regarding the number of cells composing tissues, the number of cells in the fascicular cambium (FC) and xylem (Xy) significantly increased due to prolonged exposure to hypergravity both in the apical and basal part (Fig. 7c, d). In addition, total number of cells in the internode of the primary inflorescence significantly increased under hypergravity in the basal part (Fig. 7d). Cell size significantly increased due to prolonged exposure to hypergravity in the epidermis (Ep), starch sheath (SS), phloem plus phloem cap (Ph+PhC), and fascicular cambium (FC) in the basal part (Fig. 7f) while that significantly increased only in the starch sheath (SS) and phloem plus phloem cap (Ph+PhC) in the apical part (Fig. 7e).

## Discussion

Length of the Arabidopsis primary inflorescence significantly decreased (Fig. 1a and Fig. 2a) and dry mass of the primary inflorescence per unit length significantly increased due to 10-day exposure to hypergravity (Fig. 2a, b), supporting our previous results obtained using the Arabidopsis primary inflorescence treated with 300 × *g* for 1 d (Nakabayashi et al. 2006; Tamaoki et al. 2006). Percent decrease of mean length of the primary inflorescence is 13 % in the case of 300 × *g* for 1 d when compared to 1 × *g* control (Tamaoki et al. 2006) while it is 31 % in the case of 8 × *g* for 10 d when compared to 1 × *g* control in the present study (Fig. 8a). Percent increase of mean dry mass of the primary inflorescence is 20 % in the case of 300 × *g* for 1 d when compared to 1 × *g* control (Tamaoki et al. 2006) while it is 71 % in the case of 8 × *g* for 10 d when compared to 1 × *g* control in the present study (Fig. 8b). In the present study, the peduncle could not stand the mass of inflorescence under 8 × *g* and, thereby, could not keep standing in an upright position during the entire 10 d period of exposure to 8 × *g*, which means that the peduncle had not necessarily born load of the inflorescence during this entire period. Nevertheless, effects of exposure to 8 × *g* for 10 d in length and dry mass of the primary inflorescence was stronger than to 300 × *g* for 1 d. This indicates that difference in the duration is more important than that of the gravity acceleration for the hypergravity effects on the inflorescence growth and directions of the gravitational accelerations do not affect the hypergravity effects on the inflorescence growth.

**Fig. 8.**
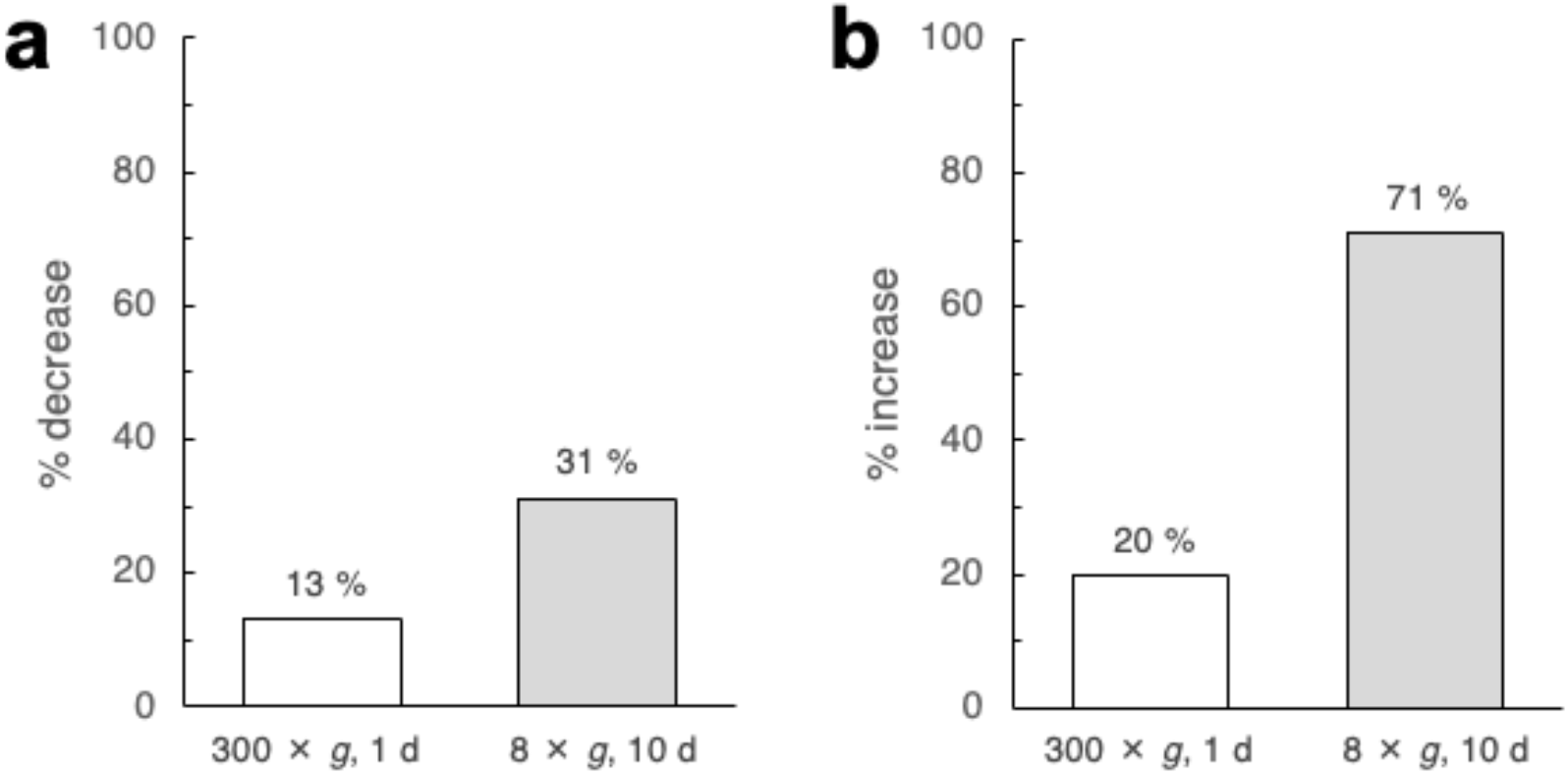
Comparisons of the effects of hypergravity exposure on the length (**a**) and the dry mass (**b**) of the main axis of the Arabidopsis primary inflorescence between the case of 300 × *g* for 1 d (Tamaoki et al. 2006) and the case of 8 × *g* for 10 d (the present study). Percent decrease of mean length of the primary inflorescence (**a**) and percent increase of mean dry mass of the primary inflorescence (**b**) when compared to 1 × *g* control.

In the present study, cross sectional area of the internode of the primary inflorescence significantly increased due to prolonged exposure to hypergravity in the basal part (Fig. 7b), which is consistent with our previous results (Nakabayashi et al. 2006) as well as the results obtained by treating *B. rapa* L. with 4 × *g* for 16 d (Allen et al. 2009). These effects, which are known as gravity resistance responses of the stem (Hoson and Soga 2003; Soga 2013), become more apparently shown in the basal internode (Fig. 3) in the present study by prolonged exposure to hypergravity.

Regarding effects of hypergravity on lignin deposition, enhancement of autofluorescence of xylem by hypergravity has been reported previously (Tamaoki et al. 2006). Therefore, we first performed anatomical observation of the main axis of the primary inflorescence and found that interfascicular fibers were developed in the basal part, which was different from our previous observation of plants after 3 days after the treatment with 24-h of 10 × *g* or 1× *g* gravity of Stage 5.10 plants. Fluorescence microscopy showed that blue autofluorescence of lignin appeared stronger at each position in the main axis of the primary inflorescence treated with hypergravity (Fig. 4), indicating that lignin deposition in xylem elements and interfascicular fibers is enhanced under hypergravity. An effect of hypergravity on formation of the fiber is newly evaluated in the present study due to prolonged exposure to hypergravity.

By performing quantification of ABS-lignin, significant increase of the lignin content in the main axis of the primary inflorescence by prolonged exposure to hypergravity was demonstrated, confirming our previous results (Tamaoki et al. 2006). Percentage increase of the mean lignin content in the main axis of the primary inflorescence by exposure to hypergravity was approximately 28 to 65 % by the exposure to 300 × *g* for 1 d when compared to 1 × *g* control (Tamaoki et al. 2006) and 39.7 % by the exposure to 8 × *g* for 10 d when compared to 1 × *g* control in the present study, which are comparable. On the other hand, content of ABS-lignin did not increase by the exposure to 4 × *g* for 16 d in *B. rapa* L. (Allen et al. 2009). Such a discrepancy in the responses to gravitational accelerations between these studies might attribute to a possibility that the threshold of this response is between 4 × *g* and 8 × *g*. Or, it might attribute to another possibility that the differences of sensitivity of plant species to gravitational acceleration. Because body size of Arabidopsis plant is larger than that of *B. rapa* L. ‘Wisconsin Fast Plant’, it is possible that the former is more sensitive to gravitational acceleration. Promotion of lignin deposition under hypergravity is suggested to be due to the enhanced activity of class II peroxidase (ATPA2) that is responsible for lignin polymerisation (Tamaoki et al. 2009), which is possibly mediated through increase of endogenous auxin level (Tamaoki et al. 2011).

In the rosette leaves of Arabidopsis, however, significant increase of the lignin content was not observed when the plants at the Stage 5.10 were treated with hypergravity (Fig. 6b), also supporting our previous results (Tamaoki et al. 2006). No significant increase was observed in lignin content in the leaf grown under microgravity in *Triticum aestivum* L. (Stutte et al. 2006). Nevertheless, lignin content slightly but significantly increased due to prolonged exposure to hypergravity in the rosette leaves of Arabidopsis when the plants at the Stage 1.02 were treated (Fig. 6c). We previously reported that rosette leaves of Arabidopsis spread obliquely upward away from the stainless plate under microgravity, while those were in contact with the plate either under space 1 × *g* and ground 1 × *g* conditions (Karahara et al. 2020). These pieces of evidence together indicate that rosette leaves of Arabidopsis also have capability to show gravity resistance response even though it is not apparent under 1 × *g* conditions on Earth.

Regarding effects of prolonged hypergravity on the stem at the level of tissues, cross-sectional area of the tissue significantly increased due to prolonged exposure to hypergravity in most type of the tissues except for interfascicular fibers (IFs) in the basal part than the apical part (Fig. 7a, b). This result that the effect of hypergravity was more prominent in the basal part indicates that the effect of hypergravity is more pronounced in the basal part than the upper part. This phenomenon is possibly because ages of cells are older in the basal internode of the primary inflorescence than the upper part, which means that the basal part is exposed to hypergravity longer than the other. This phenomenon appears reasonable considering that the function of the basal internode of the primary inflorescence is to bear load of the upper part and thus the basal internode is expected to be stronger than the upper part. On the other hand, Fig. 6 indicates that the timing of exposure to hypergravity is more important for the hypergravity exposure to take effect.

Interestingly, promotive effects of hypergravity were not observed for cell size and cell number in interfascicular fibers unlike other tissues, which was unexpected because fibers are considered to be responsible for mechanical support of the stem besides xylem. The reason for absence of promotive effects of hypergravity in interfascicular fibers might be because they developed later than other tissues and, therefore, the length of the period of exposure to hypergravity for them is shorter than for other tissues. It is necessary to examine the effect of much longer period of exposure to hypergravity on the fiber formation to confirm the absence of promotive effects of hypergravity in interfascicular fibers in the future.

Regarding effects of hypergravity on the number of cells in tissues, it was previously shown that neither of that in the pith, cortex, nor vascular tissue significantly increased in the stem of *B. rapa* L. (Allen et al. 2009). In the present study, however, only the number of cells in fascicular cambium (FC) and xylem (Xy) significantly increased due to prolonged exposure to hypergravity both in the apical and basal internodes of the main axis of the Arabidopsis primary inflorescence (Fig. 7c, d). Regarding counting xylem elements, protoxylem and metaxylem were not separately counted in the present study. However, it is considered that the number of metaxylem elements are increased due to hypergravity exposure because our previous study demonstrated that hypergravity at 300 g for 24 h did not increase the number of protoxylem elements but of metaxylem elements (Nakabayashi et al. 2006). These results indicate a possibility that hypergravity stimulates procambium activity to produce xylem element more than phloem element in Arabidopsis. Similar to the difference in the responses of the enhancement of lignin deposition to gravitational accelerations mentioned above, the difference in the responses of the enhancement of cell production in these tissues to gravitational accelerations between these studies might attribute again to the differences in plant species and/or a difference in threshold of gravitational acceleration in this response.

On the other hand, regarding the size of cells in tissues, exposure to hypergravity increased it significantly in the pith, cortex, and vascular tissue in the stem of *B. rapa* L. (Allen et al. 2009). Exposure to hypergravity also increased the size of cells in not all but many types of tissues analyzed in the basal internode of the Arabidopsis primary inflorescence (Fig. 7e, f). Increase in the size of cells in many types of tissues in the stem appears to mainly contribute to the stem thickening under hypergravity.

Regarding molecular mechanisms involved in gravity resistance responses, Ca^2+^-permeable mechanosensitive ion channels, MCAs (mid1-complementing activity proteins) are suggested to be involved in the perception of gravity signals in plants (Nakano et al. 2021) and responsible for resistance to hypergravity (Hattori et al. 2020). Because increases in cross-sectional area of metaxylem elements as well as lignin deposition in the secondary cell wall fraction by hypergravity treatment were suppressed by a blocker of mechanoreceptors, gadolinium chloride (Nakabayashi et al. 2006; Tamaoki et al. 2006), MCAs might also be involved in these responses. Gadolinium chloride, however, did not suppress promoting effect of hypergravity on an increase in the number of metaxylems, indicating that activity of procambium which determines the number of metaxylem elements is not controlled by gadolinium-sensitive mechanoreceptors (Nakabayashi et al. 2006). Factors, such as WOX4 and cytokinin signaling that are suggested to regulate procambial activity and such as HD-ZIP III that is indicated to play a role in metaxylem differentiation from procambium (Ohashi-Ito and Fukuda 2010), might be affected by hypergravity exposure. Further studies are needed to understand how procambium activity is regulated in response to hypergravity.

Cultivation of plants in space habitats is essential as a component of bioregenerative life support system for long-term exploration, utilization and development of space. It has already been demonstrated that Earth’s gravity is not absolutely necessary for plants to complete their entire life cycle by performing ‘seed to seed experiments’ in space using plant species such as Arabidopsis (Karahara et al. 2020; Link et al. 2014; Link et al. 2003; Merkys and Laurinavicius 1983; Musgrave and Kuang 2003), *Brassica rapa* L. (Musgrave et al. 2005; Musgrave et al. 2000). However, specific effects of altered gravity conditions on morphological and physiological phenomena at each developmental stage of plant’s life cycle are still largely unknown and thus examinations of these effects are important to understand and to maximize adaptive capability of plants to altered environments for manned space exploration in the future. Long-term hypergravity experiment using a commercially-available centrifuge equipped with a lighting system developed in the present study will facilitate such examinations and contribute in this area of research.

## Conclusions

We have successfully performed a lab-based long-term hypergravity experiment using a commercially-available centrifuge equipped with a lighting system. We analyzed the effect of long-term hypergravity environment on lignin deposition and on tissue anatomy in detail by comparing the effects between the positions near the apex and the base of the Arabidopsis primary inflorescence. As a result, lignin content increased in the rosette leaves as well as in the main axis of the primary inflorescence when they were treated with long-term hypergravity during their growth. Cross-sectional area of the main axis of the primary inflorescence increased under long-term hypergravity by increasing cell size in many of the tissues and the cell number in vascular tissues, such as, fascicular cambium and xylem, composing the stem. From these results it is suggested that the main axis of the Arabidopsis primary inflorescence is strengthened through changes in its morphological characteristics as well as lignin deposition under long-term hypergravity conditions.

## Acknowledgements

This work was supported fully by JSPS KAKENHI Grant (Nos. 21570064, 24620003, 15K11914) (to I. K.) and partly supported by 2022 Front loading research grant funded by Japan Aerospace Exploration Agency (JAXA) and Institute of Space and Astronautical Science (ISAS) Expert Committee for Space Environment Utilization Science.

## Conflict of interest

The authors declare no conflict of interest.

## Notes

### Competing Interest Statement

The authors have declared no competing interest.

